# MADAME: a user-friendly bioinformatic tool for data and metadata retrieval in microbiome research

**DOI:** 10.1101/2023.10.14.562335

**Authors:** Sara Fumagalli, Giulia Soletta, Giulia Agostinetto, Manuel Striani, Massimo Labra, Maurizio Casiraghi, Antonia Bruno

**Affiliations:** Biotechnology and Biosciences Department, University of Milano-Bicocca, Milan, Italy; Computer Science Institute DISIT, University of Piemonte Orientale, Alessandria, Italy

## Abstract

Microbiome research advancements have provided countless insights. Despite the massive amount of data currently stored in public repositories, these resources remain vastly underutilized due to the intricacy of data and metadata retrieval from these databases. However, leveraging data-driven approaches is crucial for microbiome research progress by overcoming variations between studies and identifying generalizable trends.

We designed the open-access and user-friendly bioinformatic tool MADAME (MetADAta MicrobiomE) to streamline the data and metadata retrieval process. MADAME addresses the challenges posed by the public repositories’ current limitations, allowing users to retrieve publications associated with the accession codes of interest. Additionally, MADAME allows users to visually explore retrieved results through the generation of a comprehensive report with plots and statistics. These unique features of MADAME let users maximize their time and resources, enabling them to assess metadata suitability before pursuing data download. To showcase its diverse functionalities, we recreate several scenarios to meet the diverse requirements that researchers may have.

## 1 Introduction

In recent years, the spread of microbiome-related studies has contributed to an exponential increase in genomics data availability. These rich datasets are a fundamental resource for meta-analyses and secondary analyses, leading to novel discoveries, data reuse approaches and data mining applications. Indeed, data-driven approaches can expand our knowledge and enhance our scientific investigation capacity in many fields of research, ranging from microbial ecology, forensics, and medicine.

Combining diverse microbiome data in meta-analyses boosts sample sizes, bolstering statistical power. Indeed, microbiome study results fluctuate due to diverse factors. Meta-analyses minimize this variation, providing a more precise estimation of microbiome effects. In this way, robust, generalizable microbiome-health relationships become identifiable (Broderick et al. 2023). Moreover, meta-analyses unveil elusive trends and associations in combined data, facilitating links recognition between microbial groups or compositions and health conditions. Often microbiome effects are subtle, but biologically significant: extending the analysis to a wider cohort can unveil such effects. Last but not least, meta-analyses offer insights for future studies, guiding research and aiding mechanistic understanding and clinical applications.

Nevertheless, there is a striking incongruence between the impressive amount of data produced and the outcome in terms of products from microbiome research both in environmental sciences as well as in the medical field is still limited. In January 2022, only 8000 microbiome-related patents were registered worldwide (Cernava et al. 2022, Espacenet – patent search). This incongruence can be explained by multiple reasons. One clear evidence is that research efforts are scattered in independent showcases, with not always shared associated metadata. Most of the well-known repositories used for microbiome data, such as MG-RAST (Meyer et al. 2008), ENA (Leinonen et al. 2011) and SRA/NCBI (Kodama et al. 2012) are all based on the the MIMARKS (Minimum Information about a MARKer gene Sequence) and MIxS (minimum information about any (X) sequence) checklist for reporting information about sequencing studies (Yilmaz et al. 2011), but it is not always effective. The varying needs of different groups working with microbiome-related data cause the absence of universally accepted metadata requirements.

A second issue, linked to the previous one, regards the common ontologies. The utilization of common ontologies, or precise technical terminology to delineate the host organism or environment, is pivotal for facilitating the reusability of metadata. Ontologies constitute hierarchies of meticulously defined and standardized lexicons, interconnected through logical relationships (Bodenreider and Stevens 2006). Their principal objective is to enhance the searchability, comparability, and machine-readability of metadata. Nevertheless, researchers frequently express dissatisfaction regarding the lax adherence to established common ontologies. Striking the balance between comprehensive information inclusion and enabling data retrieval via straightforward search queries is imperative. Consequently, several researchers opt to furnish only the bare minimum of requisite data, or employ inaccurate terminology when describing their experimental data.

These aspects hinder the aggregation of findings through integrative analyses. What we know is that standards are the key to a better usage of omics data for integrative microbiome analysis (Cernava et al. 2022) and, at the same time, users require a system that should be easy to use, clearly structured in a hierarchical way, and should be compatible with existing data repositories.

Regarding nucleotide sequence data storage, three are the main databases: the Sequence Read Archive (SRA) maintained by the National Center for Biotechnology Information (NCBI) (Kodama et al. 2012), the European Nucleotide Archive (ENA) operated by the European Bioinformatics Institute (EBI) (Leinonen et al. 2011), and the Sequence Read Archive (DRA) administered by the DNA Data Bank of Japan (DDBJ) (Mashima et al. 2017). These three databases are effectively united under the umbrella of the International Nucleotide Sequence Database Collaboration (INSDC), ensuring continuous synchronization to facilitate seamless data sharing among them (Cochrane et al. 2016).

To promote data discovery and reuse in the microbiome field, and allow for broader dissemination of knowledge and compliance for both humans and machines, data and associated metadata need to be Findable, Accessible, Interoperable, and Reusable (FAIR). FAIR principles are supported within the National Microbiome Data Collaborative and FAIR Microbiome community (FAIR Microbiome). Since data and metadata retrieval in an effective and user-friendly process still represents an obstacle for most researchers, several tools have been developed to successfully address these challenges. However, to our knowledge, tools providing such tasks do not fully cover the entire process from metadata retrieval to data download, specializing in one or few steps rather than offering a comprehensive solution to manage the lack of metadata standardization. Thus, existing tools preclude essential steps like assessment, curation, and eventually enrichment of metadata, before the actual data download. For example, some tools, like NCBImeta (Eaton 2020) and ffq (Gálvez-Merchán et al. 2023), exclusively focus on metadata retrieval, while others, including pysradb (Choudhary 2019), q2-fondue (Ziemski et al. 2022) and grabseqs (Taylor, Abbas and Bushman 2020), specialize in both metadata and data fetching and download. In particular, pysradb and q2-fondue automate metadata download from SRA, and NCBImeta extracts metadata from NCBI databases suite. On the other hand, ffq enables fetching metadata from multiple datasets (SRA, GEO, ENA, DDBJ, and ENCODE (ENCODE Project Consortium 2012, Davis et al. 2018)), as does grabseqs from SRA, MG-RAST (Meyer et al. 2008) and iMicrobe (Youens-Clark et al. 2019). MADAME is, however, the sole tool specialized in retrieving both data and metadata from ENA, the European node within INSDC. The ENA Browser is frequented by over 40,000 visitors each month and its RESTful APIs are extensively utilized for data searching and retrieval (Burgin et al. 2023). Indeed, these APIs also constitute a fundamental component of the tool. Furthermore, MADAME allows the retrieval of publications and enables the assessment of these outcomes through report generation, proving unique features among metadata and data downloading tools.

MADAME is a user-friendly, open-access bioinformatic tool that facilitates data and metadata retrieval and download from the European Nucleotide Archive (ENA). Consisting of four distinct modules, MADAME addresses the challenge posed by the absence of metadata standardization and the limited adherence to common ontology. Indeed, in addition to data and metadata retrieval, MADAME allows users to fetch the associated publications and generate comprehensive reports, including plots and statistics. This easy-to-use framework downloads data in a standardized format, according to FAIR principles, to improve the reproducibility of the analyses.

## 2 Materials and methods

MADAME is an open-source bioinformatic tool designed to streamline and automate data and metadata retrieval from the ENA database. Developed in Python 3.11 and compatible with both Linux and Windows, MADAME does not require specialized bioinformatic or programming skills: its user-friendly command-line interface is designed to guide users through the pipeline. Upon initiation, MADAME prompts users to choose between creating a new session or continuing with an existing one. All downloaded files are conveniently organized within project folders, and stored in the session directory. After choosing a session, the tool offers four main modules: (1) Metadata Retrieval, (2) Publication Retrieval, (3) Report Generation, and (4) Data Retrieval. The following sections provide a detailed description of each module.

### 2.1 Metadata Retrieval module

The core of MADAME is the module “Metadata Retrieval”, as it allows to retrieve the merged experiment metadata table that all the other modules take as input (Fig. 1). First, a list of accession IDs (unique identifiers of ENA records (Accession Numbers)) is needed. The module facilitates list retrieval through (i) a text-based query but also allows users to (ii) input a list directly or (iii) import one from a TSV or CSV file. The manual query is performed through the ENA Browser Data API (Swagger UI) after selecting the type of accession to search for: BioProject, study, experiment, sample, or run IDs. Alternatively, the user can digit a list of accession IDs, separated by comma. By matching against regex patterns, all the accession types mentioned earlier are accepted and automatically identified, so users can input mixed types as well. Additionally, we included the possibility to enter accession IDs as ranges (e.g. ERR6841562-ERR6841570) to enhance user-friendliness. Finally, the user can import a list of accessions from a TSV or CSV file by simply specifying the file path.

**Figure 1.**
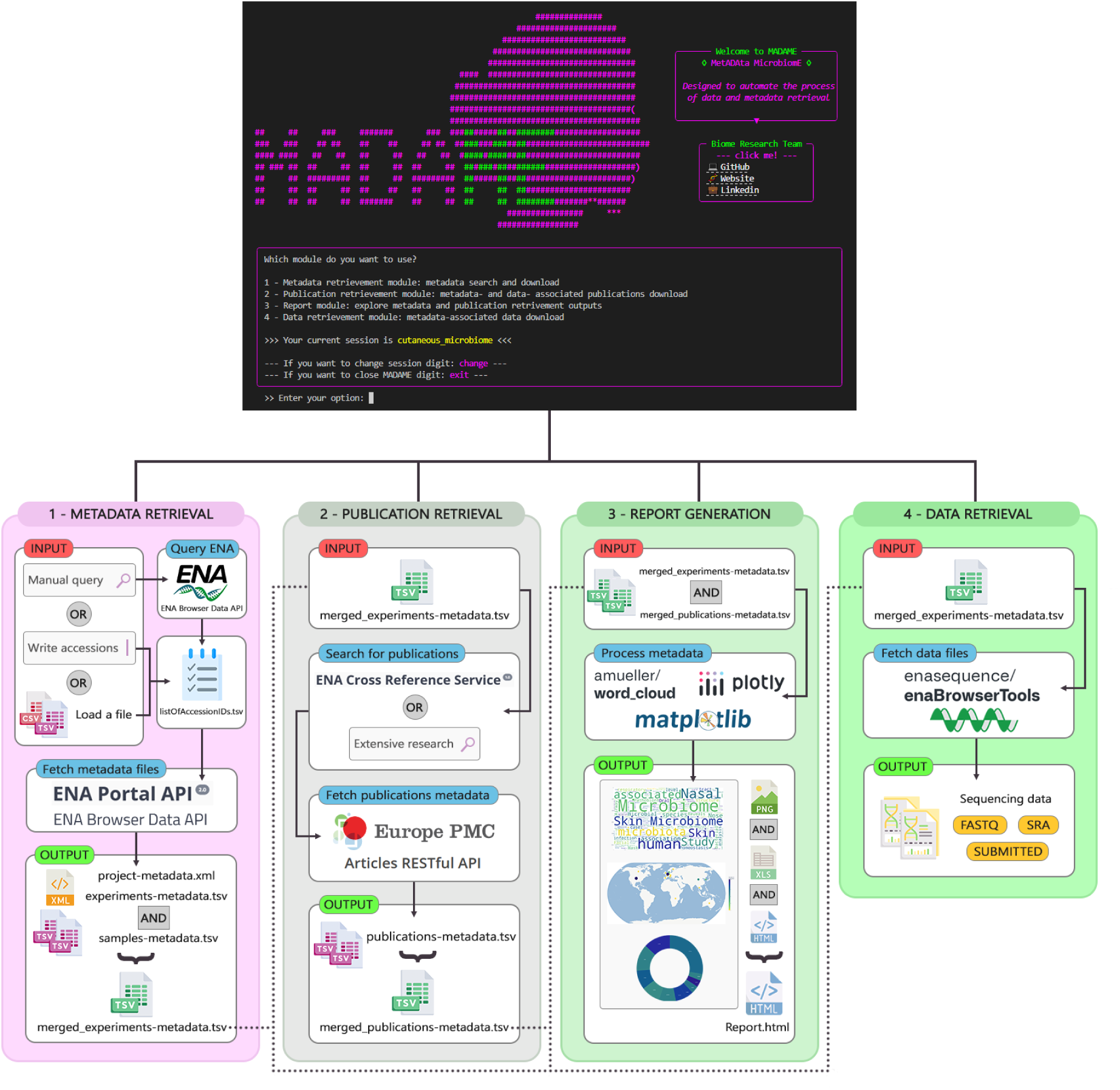
An overview of MADAME pipeline: module 1 searches and downloads sequences metadata; module 2 searches and downloads publications metadata; module 3 processes metadata and generates the report; module 4 downloads the sequencing data

In all three instances, calling the ENA Browser Data API permits the printout of accessions and their brief descriptions on the screen using the TSV response. After verifying the availability of metadata, results are printed on the screen and the “listOfAccessionIDs.tsv” file, containing only available accessions, is stored in the session folder.

Moving forward, the module prompts the user to choose between (i) downloading project and experiment metadata, along with downloading and parsing sample metadata (recommended choice), and (ii) downloading only project and experiment metadata. The project and experiment metadata are downloaded through ENA Browser Data API and saved in XML format, while the experiment metadata is retrieved from ENA Portal API (Swagger UI) and saved in TSV format. Additionally, metadata from each “experiment.xml” file is parsed and merged into a unified tabular file (“samples-metadata.tsv”) organizing each sample on a distinct row, to enhance user accessibility. The metadata parsing employs a custom parser built with the ElementTree package (xml.etree.ElementTree). All these files are organized within their respective project folders, created inside the session directory.

After downloading metadata files, the “listOfAccessionIDs.tsv” file is updated to include a second column indicating the BioProject for each accession. As a final step, experiment metadata from each project is merged into one comprehensive table file, “merged_experiments-metadata.tsv”, and saved in the session folder.

### 2.2 Publication Retrieval module

Using the “merged_experiments-metadata.tsv” file as input, the second module “Publication Retrieval” allows the retrieval of publications associated with the projects. The user is prompted to choose between (i) using the TSV file already present in the session or (ii) specifying the path to a different one. In fact, the user can customize the original file by keeping only the rows of interest, according to their preferences.

As a first step, the ENA Cross Reference Service (Swagger UI) is used to check for any publications associated with the Bioproject IDs in the session. If any publications are found, the API will supply Pubmed and/or EuropePMC identifiers. This is followed by searching for these identifiers in the Europe PMC database via its Articles RESTful API (Europe PMC), which fetches the publications metadata.

However, we have observed that the associated publications are not always indicated on the ENA website. In such instances, an extensive search is conducted: for each project, MADAME extracts a list of all the accessions from the experiment metadata and queries them on the Articles RESTful API. The API returns the metadata of each publication that includes at least one of the searched accessions in its body.

The API’s XML outputs are parsed into a tabular TSV file saved in the project folder, with one publication per row, making metadata easily accessible. If available, additional columns with download links (not explicitly present in the XML) are included: (i) Full Text XML, (ii) Supplementaries, (iii) TGZ Package from NCBI (FTP Service). Parsing is performed using an ElementTree-based custom parser.

As a final step, publication metadata from each project is merged into one comprehensive table file, “merged_publications-metadata.tsv”, and saved in the session folder.

### 2.3 Report Generation module

The third module “Report Generation” takes the “merged_experiments-metadata.tsv” and “merged_publications-metadata.tsv” files as inputs to generate a comprehensive report file. This informative report contains representative graphs and statistics, enabling users to interactively explore metadata and make informed decisions on data download, leading to significant time and resource savings. To initiate this module, the “merged_experiments_metadata.tsv” file is mandatory, while the “merged_publications-metadata.tsv” file is optional. This flexibility recognizes that users may prefer to focus solely on project and experiment metadata without the need for fetching publication metadata.

The user is prompted to choose between (i) using the TSV files already present in the session or (ii) specifying the path to a different one. MADAME initiates by identifying the TSV files within the folder and using the retrieved information to generate outputs. The primary output is an interactive HTML-format report, enabling users to explore plots by hovering over their elements, selecting specific elements of interest, and zooming into details. Furthermore, MADAME creates a folder named “Report_images” including individual HTML and PNG format plots, along with tables in XLSX format.

The plots derived from the project and experiment metadata portray information regarding the sequencing experiment (The ENA Metadata Model). This includes technical details about the library and sequencing instrument, along with biological information such as the scientific names of the collected samples. When the “merged_publications-metadata.tsv” file is provided, MADAME generates additional plots that draw from the information contained in this file, like details about publications titles and authors affiliation countries.

### 2.4 Data Retrieval module

With the fourth module “Data Retrieval”, the user has the flexibility to download sequencing data for entire projects or specific subsets, depending on the initial query or provided accessions, as well as their preferences for customizing the “merged_experiments-metadata.tsv” file.

Multithread technology provides realistic problem-solving strategies for multitask programming. In this case, the metadata download task was implemented with a multithreads mechanism (Gorelick and Ozsvald 2020) with asynchronous calls. It was observed that in contrast to the previous single-thread implementation of the “Metadata retrieval module”, the parallelized download achieved - on average - an improvement of about 20% on time complexity. This rate of improvement was achieved through different experimental runs of downloads, taking into account different factors such as number of the running processes on the CPU, network latency, and Round-trip time (RTT) from client MADAME to remote servers.

First, the user is prompted to choose between (i) using the TSV file already present in the session or (ii) specifying the path to a different one. Next, the user will be asked to indicate their format preference for the download. The available options include (i) fastq, (ii) sra, or (iii) submitted, which is the data’s original format upon submission to the ENA database. The download is performed by employing the enaBrowserTools script package (enaBrowserTools 2023), developed by ENA. Data files are organized within their respective project folders, inside the session directory.

### 2.5 System design and functionality

Systems design is essential in information technology and information systems (IS). Systems design is an approach that emerged around the 1970s that addresses business requirements and technical issues related to software development (Siau et al. 2022). System design is typically considered a set of procedures to define the various system elements and components that adhere to a particular set of requirements (Valacich and George 2017). Conversely, system analysis comprises a process of immersing and understanding the users’ experience with a specific system in order to improve or design a new approach based on the specific requirements (Dennis, Wixom and Tegarden 2020). Therefore, system analysis and system design are two topics merged into one IS course (Kohli and Gupta 2002). In essence, system design and system analysis are complementary processes. System analysis aims to understand user experience and input, which can inform the design process, unlike system design, which focuses on developing a design for the system. To ensure that the system meets requirements and succeeds in meeting user demands, a mix of these two approaches is essential. Consequently, the system development process must include both system design and system analysis.

System modeling and analysis is characterized by two main approaches: (i) the traditional structured method and (ii) the object-oriented method (Harris et al. 2006). The traditional method is divided into two phases: the analysis of the system and its design through the use of a series of diagrams such as data flow diagrams and entity-relationship diagrams. The second is commonly understood to be data-centric and uses the Unified Modelling Language (UML), which is a set of entities, i.e. “classes” that encapsulate the data known as “attributes” and the processes “methods” related to each entity (Siau et al. 2022).

The UML is a language proposed in the 1990s and adopted in practice to model software requirements (Bucchiarone et al. 2020). The UML is also seen as a collection of different approaches, such as object-oriented notations known as object-oriented design, object modeling technique and object-oriented software engineering (Gomaa 2006). The UML has been widely adopted in teaching systems analysis and design (Tanner and Scott 2015, Burton and Bruhn 2004), which combines several diagrams to represent the behaviours and characteristics of a system. Typically, a single diagram is a graphical representation of a particular part of the target system. Ultimately, a system model comprises several diagrams to illustrate the target design.

In order to describe in a formal way the architecture and the different functionalities offered by MADAME software, we use Visual Paradigm software (described in Section 2.5.1). In the following sections we will illustrate two types of UML diagrams: the use-case diagram (see Section 2.5.2) and the class-diagram (see Section 2.5.3). In particular, the first is to highlight the dependencies between the modules and software components, and the second is to describe the classes and the inclusion relationships between them.

#### 2.5.1 Visual Paradigm and UML

Use-case and class diagrams in Visual Paradigm (Visual Paradigm Selected Users List, Free UML, BPMN and Agile Tutorials, Component Diagram - UML 2 Diagrams - UML Modeling Tool) or other modeling tools help visualize the high-level structure of a software system by illustrating how different components interact and depend on each other. These diagrams are particularly useful for designing and understanding complex systems, identifying relationships, and managing dependencies between various software components.

Visual Paradigm is a modeling tool supporting UML 2 that allows users to create different types of diagrams, including class diagrams, component diagrams, and use case diagrams, among others. It supports integration with development environments like Eclipse, IntelliJ, and Visual Studio through its API, which is implemented using Java. Overall, Visual Paradigm is a versatile tool used by various companies for designing, modeling, and managing software systems, offering a range of diagram types and plugin capabilities to cater to different needs.

A component diagram assists a designer in modeling the architecture of software components and their dependencies within an object-oriented software system (Rumpe 2016).

#### 2.5.2 Use-case diagram

A UML (Unified Modeling Language) use-case diagram is a graphical representation of the functional requirements of a system from the perspective of its users. It helps to visualize the interactions between different actors (users or external systems) and the system itself. Use case diagrams consist of actors, use cases, and their relationships. Here is how the different components are represented:

##### Actor

An actor represents a user or an external system that interacts with the system you’re modeling. An actor can be a person, another software system, or even a hardware device. Actors are depicted as stick figures on the diagram. In Figure 2, the actor represents the user (scientist/bioinformatician) using the MADAME system by interacting with its different functionalities divided into four modules, each containing the relevant use cases

**Figure 2.**
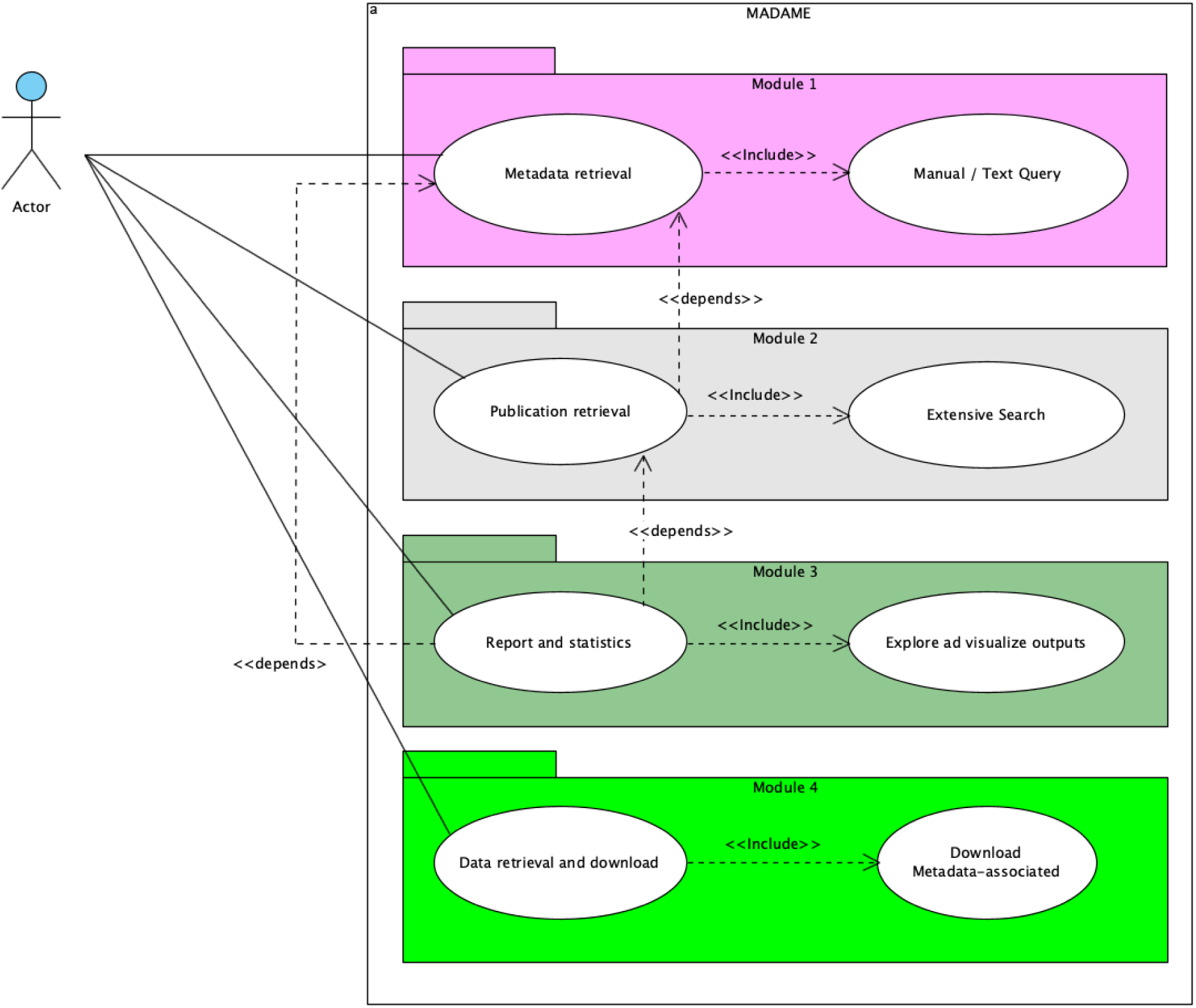
MADAME Use-case diagram

##### Use Case

A use case represents a specific functionality or behavior of the system that provides value to the actors. It describes a sequence of actions performed by the system and the actors to achieve a specific goal. Use cases are represented as ovals on the diagram.

###### Association

A solid line connecting an actor to a use case indicates that the actor interacts with or uses that particular use case. In our case, Figure 2 shows that the actor can interact with the system MADAME by using four different modules divided into different use-case diagrams.

###### Include

A dashed arrow from one use case to another indicates that the behaviour of the included use case is inserted into the behaviour of the base use case under certain conditions. In Figure 2, the use-case called “Metadata Retrieval” (defined in Module 1) includes the use-case called “Manual / Text query” (also defined in the same Module 1), because the user can use a manual sub-process for downloading projects, which requires the input of a query string. With regard to Module 2, the retrieval of publications includes the use-case extensive search. For Module 3, the actor, in order to be able to view the report and statistics, calls up a (sub)use-case to display the output on its dashboard. Finally, regarding Module 4, Data Retrieval use-case includes “Download Metadata-associated”, a sub-case allowing the download of associated metadata.

###### Dependency

A dependency is a relationship that signifies that a single or a set of model elements requires other model elements for their specification or implementation. This means that the complete semantics of the depending elements is either semantically or structurally dependent on the definition of the supplier element(s). In Figure 2, the use-case called “Publication Retrieval” (defined in Module 2) depends on the “Meta retrieval and download project” (defined in Module 1) because, in order to download projects, it will first be necessary to temporally retrieve the publications in which the project code is listed. The same for “Report Generation” (defined in Module 3), “Publication Retrieval” (defined in Module 2) and “Metadata retrieval and download project” (defined in Module 1). Figure 2 shows the UML use case diagram for the system MADAME. Finally, the use-case called “Data retrieval” (defined in Module 4) depends on “Metadata retrieval and download project” (defined in Module 1) and, for this reason, in order to retrieve data, needs to be performed after.

The system MADAME is composed of four different modules, each of which defines two different use cases; in particular,

- Module 1 is composed of “Metadata Retrieval” which includes “Manual / Text query”.
- Module 2 is composed of “Publication Retrieval” which includes “Extensive Search”
- Module 3 is composed of “Report Generation” which includes “Explore and visualize outputs”
- Module 4 is composed of “Data Retrieval and download” which includes “Download metadata-associated”

The Actor (MADAME’s user) interacts with the entire system MADAME, consisting of four modules described above. Remember that use-case diagrams are just one part of the UML and they are used to capture high-level functional requirements and interactions. They provide a quick overview of the system’s functionality and user’s interactions. More details are described in Figure 2.

#### 2.5.3 Class diagram

In the guided design of software models and systems, UML class diagrams are widely used to model the structures of a system (Atkinson and Kuhne 2003, Booch 2017). A UML class diagram is a type of diagram used in software engineering and modeling to visually represent the static structure of a system or software application in terms of classes, attributes, methods, and relationships, facilitating the understanding and design of software architectures. In a system, an entity is typically represented as a class, and inheritance or associations are represented as relationships between different entities.

Below, we will briefly describe the most important components, characteristics, and concepts used to describe the software structure of MADAME within a UML class diagram (shown in Figure 3).

**Figure 3.**
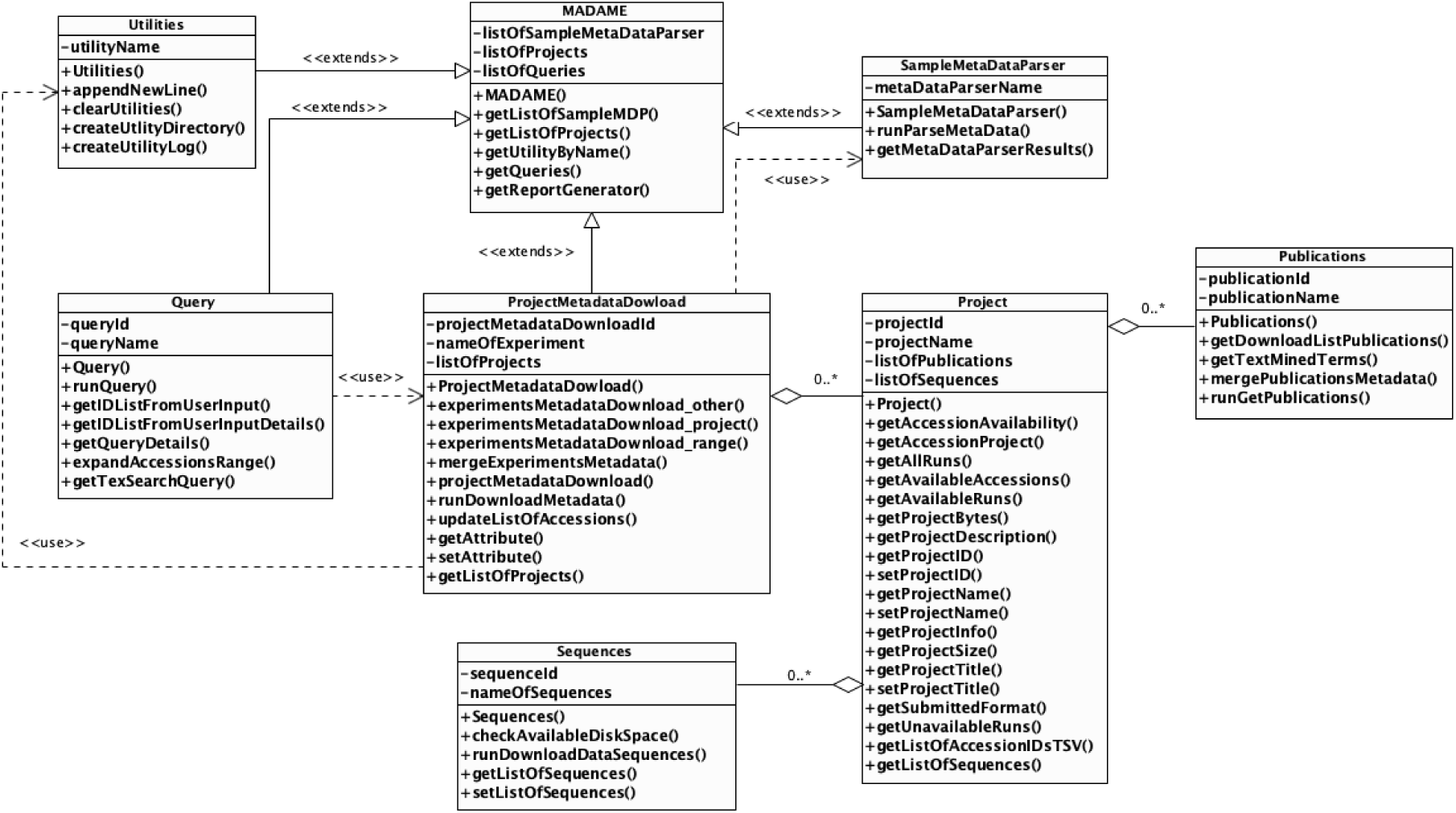
The UML class diagram of MADAME

##### Most important Classes

###### Madame

Represents The main class of the entire system MADAME, which is extended by all other classes (*Utilities*, *SampleMetaDataParser, Query,* and *ProjectMetadataDownload*). The most important method defined in this class is *getReportGenerator()* which allows to generate an informative report (Module 3) containing representative graphs and statistics. In this way, the users can interactively explore metadata and make significant decisions (see Section 2.3).

###### Project

This class represents a single project, uniquely identified by its ID and name (respectively ProjectID and projectName attributes). Each project defines within it a set of sequences (composition relation (0…*) of class Sequences) and publications (composition relation (0..*) of class *Publications*).

###### ProjectMetadataDownload

This class represents Module 1, and contains all functionalities to retrieve metadata related to the projects (see Section 2.1). This class is characterized by a set of projects (composition relation (0..*) of class *Project*). This class uses the *Query* and *SampleMetaDataParser* classes.

###### SampleMetaDataParser

This class extends the mainclass MADAME and is used by the *ProjectMetadataDownload* class to parse the project’s samples metadata.

###### Publications

This class represents the implementation of Module 2 with all methods for the retrieval of publications associated with a single project (composition relation (0..*) with class *Project* because we assume that there are projects that may not have associated publications (see Section 2.2).

###### Sequences

This class represents Module 4, where the user can download sequencing data for the entire project. (see Section 2.4). Class *Sequences* is in a composition relationship (0..*) with class *Project* because we assume that there is the possibility that a single project may not have associated sequences.

###### Query

This class represents methods for defining and executing a text-based query for the project’s metadata retrieval, (see Section 2.1). This class is used by *ProjectMetadataDownload* (relation <<USE>> with class *ProjectMetadataDownload* and extends the main class *MADAME*.

The design of the MADAME system according to the object-oriented programming paradigm, allows for an easy extension of the system and its functionality, in order to be able to add more classes, attributes, and methods that will model further developments and future extensions.

## 3 Results and Discussion

To provide a comprehensive understanding of MADAME’s capabilities, we conducted multiple tests involving different microbiome environments and various accession codes. These case studies were designed to represent commonly encountered scenarios by researchers investigating microbial communities across diverse ecosystems. By testing MADAME with a variety of retrieval and download tasks, we aim to show MADAME’s capacity to fulfill researchers’ needs through its modules’ features.

### 3.1 Input types

To illustrate the various input types accepted by MADAME, we reproduced four scenarios tailored to the diverse requirements that researchers may encounter. Each scenario takes place in the Metadata Retrieval module of MADAME, using a different input type: (i) free-text search, (ii) list of accession codes, (iii) upload of TSV or CSV format file. On October 9 2023, the retrieval and downloading of all scenarios were executed on MADAME.

#### 3.1.1 Free-text search

With its extensive surface area, the skin constitutes a complex ecosystem where microorganisms and our cells engage in continuous relationship with the dynamic external environment. A key objective in microbial ecology is to establish connections between the phylogenetic diversity of microbes associated with hosts and their functional contributions to the community. There are still significant knowledge gaps regarding the mechanisms of microbial colonization on the skin and how the skin microbiome responds to the environment. To address this, collaborative endeavors and interdisciplinary research bridging environmental and medical microbiology are imperative to aggregate knowledge and propel practical investigations (Byrd, Belkaid and Segre 2018, Callewaert, Ravard Helffer and Lebaron 2020).

In our first scenario, we aimed to investigate the skin microbiome associated with the face, while excluding the ears. We initiated the metadata retrieval process by formulating a quite intricate text query, namely “((((((((human skin) AND face) OR facial) OR cheek) OR forehead) OR glabella) OR eyelid) OR nose) AND microbiome”, focusing on projects. MADAME retrieved 46 projects, including a total of 15531 samples. By “sample” we refer to the biological sample collected, each of which corresponds to a unique file containing DNA sequences, such as a fastq file.

In our second scenario, we focused on the virome. Indeed, over the past decades, a multitude of emerging viruses originating in wildlife have been transmitted through spillover events, leading to severe outbreaks in humans. In particular, bats are renowned reservoirs of zoonotic viruses, harboring, along with primates and rodents, a higher proportion of viruses than the other mammal groups (Olival et al. 2017). Among these bat-origin pathogens, coronaviruses, filoviruses, and henipaviruses are responsible for half of the priority diseases identified by the World Health Organization as the greatest public health risk (Prioritizing diseases for research and development in emergency contexts), resulting in dramatic human and economic costs. For this reason, in recent years, there has been a commitment to transition the investigation of bat-borne diseases from a reactive approach to a proactive strategy to prevent further spillovers, by integrating virome high-throughput sequencing with other unbiased strategies (Letko et al. 2020).

Then, we performed metadata retrieval by initiating a search for studies using the free text query “bat virome”. MADAME retrieved a total of 12 studies, comprising 785 samples.

#### 3.1.2 List of accession codes

With rigorous cleanliness standards, frequent antibiotic use, and the constant introduction of new pathogens, hospitals represent an extreme environment where multidrug-resistant microorganisms thrive and can be transmitted to patients. Constituting a significant healthcare burden and an alarming global health issue, the antibiotic resistance epidemic has focused attention on understanding the transmission dynamics between hospital microbiota and patients (Chng et al. 2020, Hernando-Amado et al. 2019).

In our third scenario, we examined BioProject PRJNA544954 (https://www.ebi.ac.uk/ena/browser/view/PRJNA544954), a pilot study centered around an Egyptian hospital microbiome, which includes a total of 9 experiments related to the microbiota associated with door knobs and 3 experiments related to bed sheets microbiota. We chose to focus solely on the door knob experiments and conducted a metadata retrieval by entering the specific codes. Thanks to MADAME’s capability to input code ranges, we were able to enter the term “SRX5904061-SRX5904068, SRX5904070”, instead of inputting the 9 individual codes. MADAME successfully retrieved all the metadata associated with these experiments.

#### 3.1.3 Upload of TSV or CSV format file

The microbial diversity present in diverse vitivinicultural regions significantly influences the unique characteristics of wine terroir. It is essential to thoroughly investigate and conserve this biodiversity, although there is potential for future manipulation through precise agricultural and oenological practices. The soil microbiome has gained considerable importance within the realm of viticulture. Recent years have seen numerous studies emphasizing the profound impact of soil microbial diversity on the distinctive terroir of wine, raising the possibility of its predictive capability for determining the geographical origin of the wine. However, achieving this necessitates a comprehensive meta-analysis, integrating microbiome data from various studies, to construct a predictive model capable of distinguishing microbial patterns and accurately predicting the geographic origin of the samples (Gobbi et al. 2022, Mezzasalma et al. 2017).

Thus, in our last scenario, we investigated BioProject PRJNA432813 (https://www.ebi.ac.uk/ena/browser/view/PRJNA432813) which examines the presence of bacteria and fungi in vineyard soil, grapes, and winery during spontaneous fermentation. Within this project, our specific focus was directed towards the investigation of fungal sequences. Due to the large amount of accession codes involved, we opted to compile a CSV format file with the 134 accession codes of the fungal runs. Then we provided MADAME with the file path location of the CSV file, enabling the tool to search for metadata associated with the accession codes contained in it. MADAME successfully retrieved the experiment and sample metadata of all the runs.

### 3.2 Publications

Retrieving publications is optional but serves as a valuable tool to validate the downloaded metadata and potentially enrich them with information from publications materials and methods or supplementary sections. This approach exemplifies MADAME’s commitment to overcoming challenges arising from the lack of metadata standardization.

To test MADAME with the high number of projects and samples retrieved with the free-text search, we decided to focus on the exploration of the first scenario concerning human skin face microbiome. Thus, we initiated the Publication Retrieval module aiming to fetch the publications associated with the projects that MADAME retrieved in the previous module. MADAME retrieved 45 publications belonging to 33 projects out of the 46 with available metadata. Noteworthy, MADAME could not retrieve publications for every project. This limitation stems from the methodology of extracting all identifiers from the downloaded metadata file and querying the Europe PMC database through its Articles RESTful API. Consequently, MADAME is capable of exclusively retrieving publications that have their full-text articles made accessible. The sole exception applies to publications linked to the BioProject on ENA, which are retrieved irrespective of their accessibility. Moreover, The highest number of publications associated with a single project was four, and that occurrence was observed for two projects, while for the majority of projects, MADAME retrieved one publication. MADAME retrieved even the publications linked to a project cited in the manuscript and/or used for the meta-analysis addressed in the publication. This is the case of the BioProject PRJNA269493 (https://www.ebi.ac.uk/ena/browser/view/PRJNA269493), where MADAME identified three publications, one of which is a meta-analysis (Shi et al. 2019).

The oldest publication dated back to 2012, whereas the most recent publications were from 2023. However, the majority of the publications were dated 2021. According to the authors’ affiliations, publications were distributed across 17 countries, with an equal distribution of ten in the USA and ten in Belgium (Fig. 4a). The worldcloud of project-associated publications titles showed a predominance of words related to the skin sampling site (“nasal”, “nose”, “facial”, “oral”), but also references to diseases (“atopic dermatitis”, “pathobiont”, “infection”). Noteworthy, also the words “mask”, “respiratory”, “airway” appeared in the wordcloud, probably due to the recent research efforts about COVID-19 pandemic (Fig. 4b).

**Figure 4.**
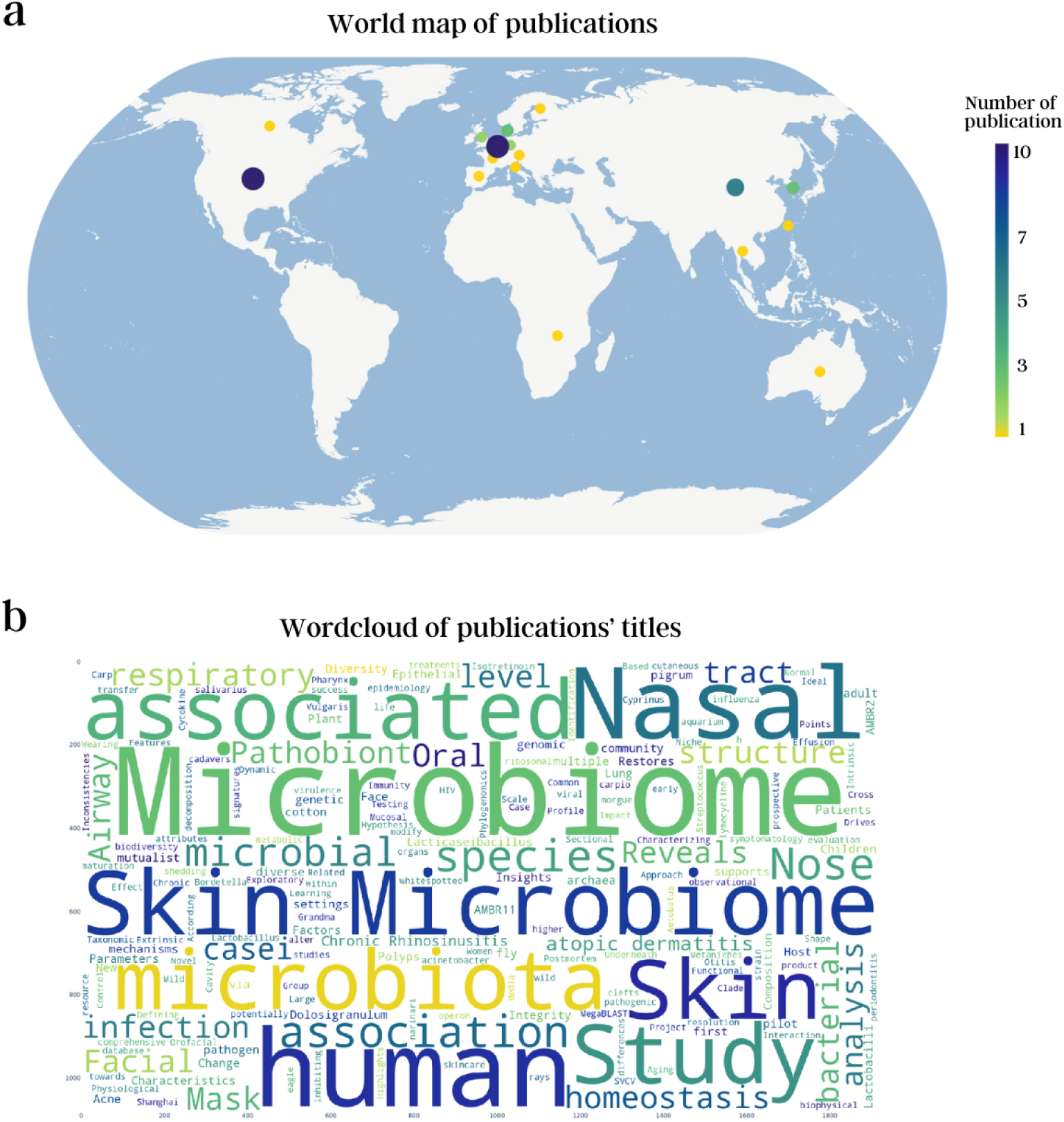
a) Word map of the publications. b) Wordcloud of the publications’ titles. All the plots were generated through Report Generation module.

### 3.3 Report

Considering the extensive amount of metadata retrieved by MADAME, encompassing both experiments and publications, we explored these data through the generation of a comprehensive report. Similar to the publications module, the report generation module is optional. Nevertheless, this distinctive aspect of MADAME enables users to visualize results swiftly yet effectively, providing a clear overview of the downloaded metadata. Indeed, the absence of standardization and limited adherence to ontologies can result in the retrieval of projects unrelated to the initial query, necessitating filtering before downloading their data. Thus, we initiated MADAME’s third module to produce a comprehensive report on projects and publications metadata. With access to the relevant publication metadata, MADAME efficiently generated supplementary plots, enabling the visualization of crucial information about both the experimental metadata (Fig. 5) and publications (Fig. 4).

**Figure 5.**
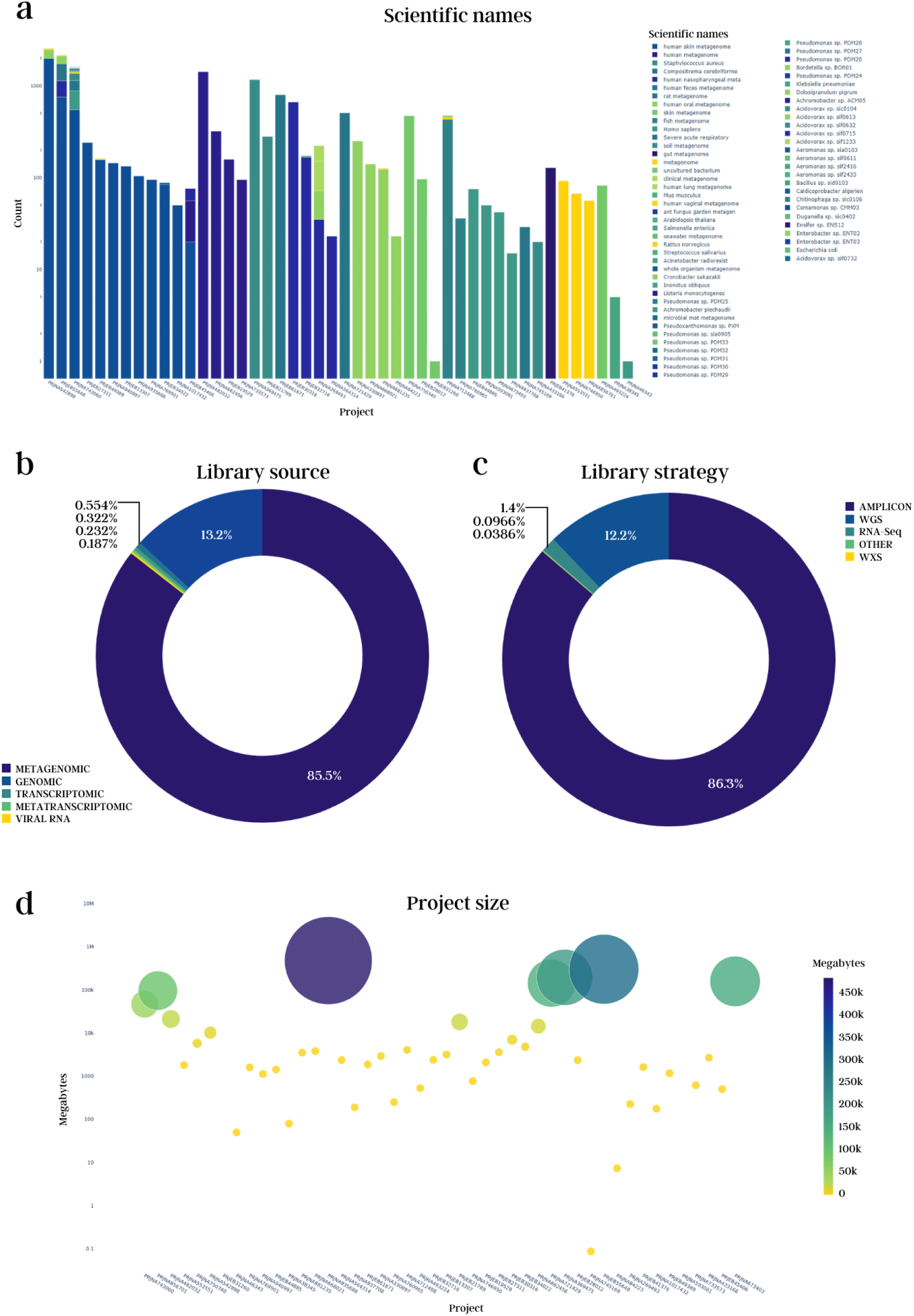
a) Bar plot of the scientific names. b) Donut chart of the library source. c) Donut chart of the library strategy. d) Scatter plot of the projects’ size. All the plots were generated through Report Generation module.

MADAME uncovered 68 scientific names associated with the 46 projects identified in our initial query into the human facial skin microbiome. Despite the median count of scientific names per project being 1, one project stands out with 48 names (PRJNA743060, https://www.ebi.ac.uk/ena/browser/view/PRJNA743060) representing the source of variability of scientific names (Fig. 5a). The three most abundant scientific names, determined by the number of associated samples, were as follows: “human skin metagenome” at 27.7%, “human metagenome” at 13.1%, and “other” at 13% which included all scientific names absent in the top ten most abundant. At lower percentages, we found scientific names that were unrelated to our initial query. Examples included human microbiome from other body sites like “human feces metagenome”, “gut microbiome”, and “human lung microbiome” but also disparate organisms or environments, such as “rat metagenome”, “fish metagenome”, “soil metagenome”, “clinical metagenome”, and “ant fungus garden metagenome”.

The metadata exhibited the highest heterogeneity when considering scientific names. This outcome can be attributed to the lack of metadata standardization and limited adherence to ontologies. This is evident in cases for the scientific name “human microbiome”. This choice reflects a broader category rather than specifying the exact body site from which the samples were obtained.

However, the main issue arises from samples with scientific names unrelated to our query. When considering scientific names linked to other organisms, like rats and fish, it is plausible that the problem originates from how we formulated our query. Indeed, the metadata retrieval, conducted through querying the ENA Browser Data API, is consequently subject to the rules of the EBI Search engine (EBI Search Documentation).

The majority of retrieved projects contained metagenomic samples, followed by genomic, transcriptomic, and metatranscriptomic samples (Fig. 5b). As expected, we focused our search on metagenomic samples by including the term “microbiome” in our query. The prevalent sequencing method was amplicon, followed by whole-genome sequencing (WGS), and RNA sequencing (RNA seq) ranked third (Fig. 5c). Lastly, Illumina was the predominant platform at 81.8%, followed by LS454.

While metadata is accessible for many projects, not all of them have downloadable data. There are several reasons for this, and one of them is the privacy of the data (Data Availability Policy). In the case of our query, two projects (PRJNA593081; https://www.ebi.ac.uk/ena/browser/view/PRJNA593081, PRJEB31260; https://www.ebi.ac.uk/ena/browser/view/PRJEB31260) had no fastq associated that we could download. Specifically, PRJNA593081 did not have publicly associated data, while PRJEB31260 had available data only in submitted format. As for the remaining 44 projects, their sizes vary significantly: the project with the largest dimensions was PRJNA935688 (https://www.ebi.ac.uk/ena/browser/view/PRJNA935688) for a total of almost 480 Gigabytes, while the smallest was the PRJNA745169 (https://www.ebi.ac.uk/ena/browser/view/PRJNA745169) project of 80 Kilobytes (Fig. 5d).

As evident, there are 46 publications on the world map (Fig. 4a), even though the publications retrieved by Madame were 45. This discrepancy arises because MADAME relies on authors’ affiliations to extract the country of origin. In this particular instance, within the publication (Noguera-Julian et al. 2017) authors from different countries have collaborated. Noteworthy, publications lacking information about the country of origin in the authorship are not displayed on the map. Even if this occurrence is infrequent, we can observe that in two retrieved publications (Lee et al. 2019) which indicated only the internal state such as ”Baltimore, MD” omitting “USA”. Furthermore, the predominance of publications in North America and Europe is evident. The issue with this distribution highlights the asymmetry in the representation of human microbiome data, as acknowledged and described in the paper. (Abdill, Adamowicz and Blekhman 2022).

In conclusion, the combination of the Report Generation module and the Publication Retrieval module proves to be an invaluable resource, successfully mitigating the observed heterogeneity that researchers may encounter when retrieving data for reanalyses.

## 4 Conclusions

As stated in the latest ENA report, its visitors download around 350,000 GB of data each month out of the 40 petabytes it archives (Burgin et al. 2023). Considering this figure represents only 1% of the total data, it is evident that efficient data reuse is still a distant goal. The current bottleneck hindering the complete utilization of data stored in public databases lies in the difficulty for most researchers in retrieving and downloading them from repositories. To address this issue, we designed MADAME (MetADAta MicrobiomE) with the aim of comprehensively handling the entire necessary process of metadata assessment and eventual enrichment, before data downloading.

In adherence to the FAIR principles (Findability, Accessibility, Interoperability, and Reusability) (FAIR Microbiome), MADAME stands as an inclusive tool, offering researchers access to necessary resources for conducting downstream analyses, even for those lacking specific bioinformatic skills. The scenarios presented in this manuscript illustrate a subset of (albeit simplified) applications for the tool. We foresee numerous additional applications for analyzing a wide range of nucleotide sequence data types, to help users in unveiling the potential hidden in data and metadata stored in public repositories. In essence, MADAME can effortlessly supply data and associated information retrieval, with the ultimate aim of contributing to microbiome research progress by facilitating multiple datasets integration.

## Funding

This research was realized within the MUSA – Multilayered Urban Sustainability Action – project, funded by the European Union – NextGenerationEU, under the National Recovery and Resilience Plan (NRRP) Mission 4 Component 2 Investment Line 1.5: Strengthening of research structures and creation of R&D “innovation ecosystems”, set up of “territorial leaders in R&D”.

This work was funded also by the National Plan for NRRP Complementary Investments (PNC, established with the decree-law 6 May 2021, n. 59, converted by law n. 101 of 2021) in the call for the funding of research initiatives for technologies and innovative trajectories in the health and care sectors (Directorial Decree n. 931 of 06-06-2022) - project n. PNC0000003 - AdvaNced Technologies for Human-centrEd Medicine (project acronym: ANTHEM). This work reflects only the authors’ views and opinions, neither the Ministry for

University and Research nor the European Commission can be considered responsible for them.

## Data availability

An introductory tutorial and additional information for the utilization of MADAME are accessible at: https://github.com/BiomeResearchTeam/MADAME

